# Direct Nuclear Delivery of Proteins on Living Plant via Partial Enzymatic Cell Wall Digestion

**DOI:** 10.1101/2024.09.03.611064

**Authors:** Qufei Gu, Nathan Ming, Xiaoyang Wei, Yalikunjiang Aizezi, Yizhong Yuan, Brian Esquivel, Zhiyong Wang

**Affiliations:** Department of Plant Biology, Carnegie Institution for Science, Stanford, California, 94305, United States

**Keywords:** green fluorescence protein, nuclear localization sequence, peptide, cell wall, partial enzymatic digestion

## Abstract

While many variations of protein delivery methods have been described, it can still be difficult or inefficient to introduce exogenous proteins into plants. A major barrier to progress is the cell wall which is primarily composed of polysaccharides and thus only permeable to small molecules. Here, we report a partial enzymatic cell wall digestion-mediated uptake method that efficiently delivers protein into the nucleus of plant cells. Such a method allowed efficient nuclear delivery of GFP proteins into Arabidopsis root cells throughout all cell layers. This study establishes that a partial enzymatic cell wall degradation could enable a myriad of plant biotechnology applications that rely on functional protein delivery into walled plant cells.

## INTRODUCTION

The delivery of exogeneous biomolecules into walled plant cells enables targeted genetic engineering of plants. While several tools exist for the delivery of nucleic acids in plants, it is still difficult or inefficient to introduce proteins into plants. Although synthetic carriers such as nanomaterials^1,7-8^ and cell penetrating peptides (CPPs)^7, 9-10^ allow proteins to cross rigid and multilayered cell wall and phospholipid bilayer, the applications of these platforms in plant genetic engineering is limited due to low reproducibility and efficiency. A well-known challenge is that these carrier-based methods often have significant platform-to-platform, batch-to-batch and even device-to-device variabilities, especially in light of growing evidence that nanoparticles (NPs) are not required for biomolecule delivery in plants but instead provide protection against degradation.^11^ Another major barrier to the use of nanotechnology is the lack of quantitative validation of successful intracellular protein delivery, which is difficult to distinguish from artifact and lytic sequestration.^8^ New methods need to be invented to make protein delivery rapid and efficient for some plant species and possible for other species that cannot yet be accomplished using existing methods.

In this technical report, we present a new approach to achieve the goal of protein delivery in walled plant cells, as evidenced by GFP signal in the nucleus that is greater than the cytoplasmic signal. This is accomplished by partially digesting the cell wall of young Arabidopsis seedlings with enzyme (PECWD, partial enzymatic cell wall digestion), allowing cargo proteins to enter the cytosol of plant cells (Fig 1a-c), in a way that is analogous to the protoplast transformation.^12^ Our assumption is the delivery of proteins in to plant cells does not need to be preceded by the complete removal of cell wall. Unlike mammalian cells, the plant cell cytosol is tightly compressed against the cell wall by the vacuole, which makes it difficult to make unambiguous imaging of cytosolic contents. Moreover, plant cells are heterogeneous in shape and have many auto-fluorescent bodies, making distinguishing signal from background noise challenging. Here we show that the addition of two flanking NLS motifs can help to localize the GFP or GFP-tagged cargos to the nucleus, producing a round and uniform object that is amenable to image analysis and provides unambiguous confirmation and comparison of successful intracellular protein delivery (Fig 1d).^2, 13^ It should be noted that higher enzyme concentration and longer digestion period did not result in larger protein (Cas9) localization in digested seedlings. Such discrepancy can be readily explained by the incomplete cell wall removal during PECWD process, which rejects large Cas9 proteins while allowing small GFP proteins to pass through. As the aim of PECWD is to improve the permeability without losing the overall integrity and stability of the cell wall structure, a size exclusion limit should be expected. Future studies that utilize lipid- and polymer-based nanoparticles to encapsulate and condense proteins can serve to address this limitation.^7, 17^ This novel approach may have broad utilities in plant transformation techniques that require the introduction of macromolecules, and may enable the tissue culture-free gene editing methods.

**Figure 1.**
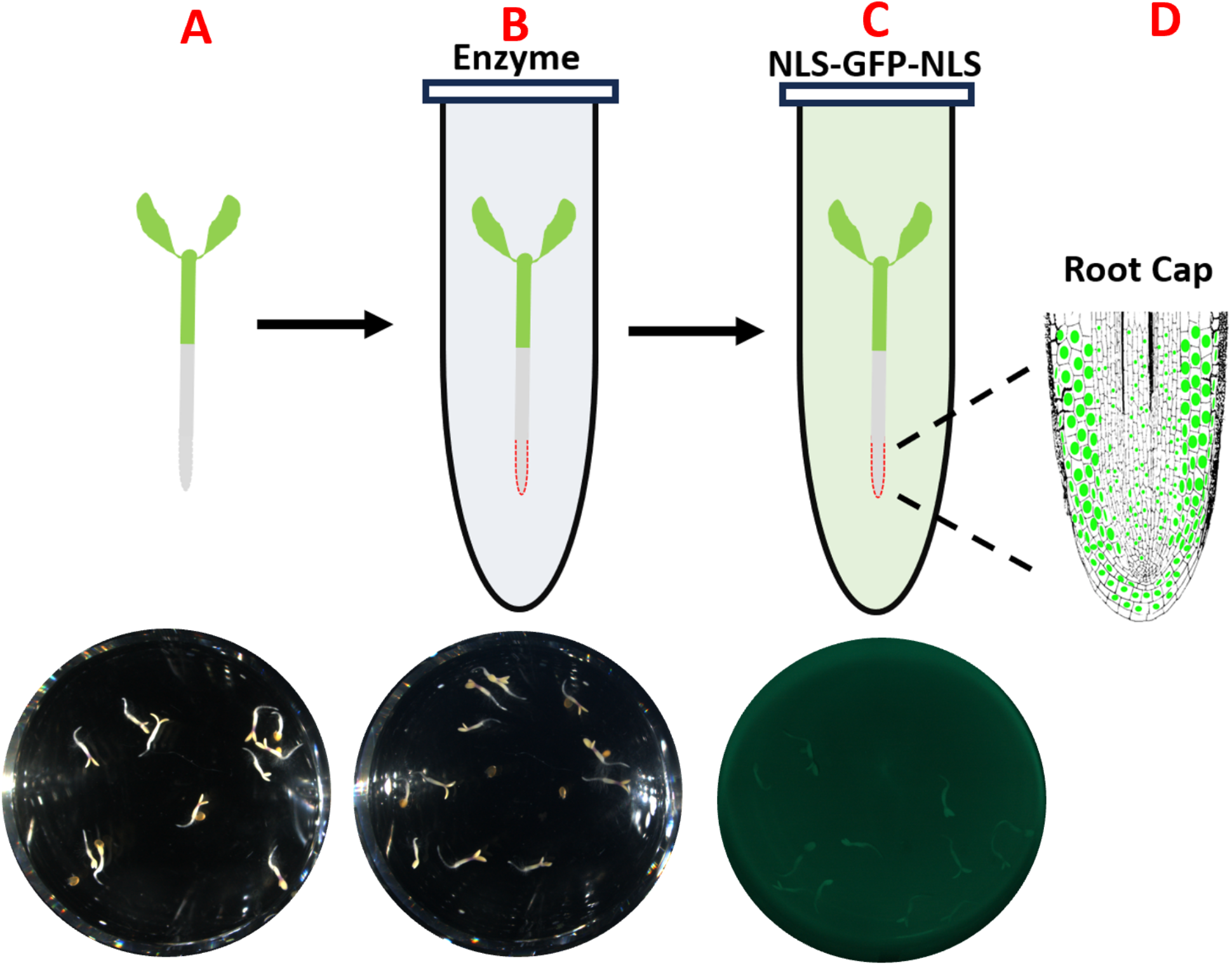
Schematic overview of nuclear protein delivery through the combination of partial cell wall digestion and nuclear localization sequence (NLS). (A) First, grow wild-type Arabidopsis seedlings in ¼ MS liquid medium under daylight or far-red light. (B) The 3-day-old seedlings are incubated in 5% hemi-cellulase solution in dark for 4 h. (C) The seedlings are then incubated in 1 mg/mL protein solution in dark for 15 h. (D) Post-incubation, seedlings are imaged on a confocal laser scan microscope to confirm the nuclear delivery.

## MATERIALS AND METHODS

### Materials

Hemi-cellulase and pectinase were purchased from Sigma-Aldrich Inc. (St. Louis, MO). Macerozyme were obtained from GoldBio Tech. Co. (St. Louis, MO). All gene fragments were synthesized by Twist Bioscience (San Francisco, CA). Phusion High-Fidelity DNA Polymerase was purchased from Thermo Fisher Scientific Inc. (Waltham, MA). Restriction enzymes and Gibson assembly kit were obtained from New England Biolabs (Ipswich, MA). *E. Coli BL21* strain harboring pET28a-Cas9-His were purchased from Addgene Inc. (Watertown, MA). Unless otherwise stated, all chemicals were obtained from Fisher Scientific Co. (Pittsburgh, PA).

### Recombinant protein expression and purification

For *in vitro* expression of GFP protein, the GFP coding sequence flanked by two SV40 NLS motifs was inserted into pET28a (+) to generate pET28a-NLS-GFP-NLS-His using Gibson Assembly. The recombinant vector was transformed into *E. Coli Top-10* strain for plasmid extraction and sequencing. Plasmids with desired sequences were then transformed into *E. Coli BL21* strain, expressed at 16°C, purified by nickel chromatography column and dialyzed in HEPES buffer (20 mM HEPES pH 7.5, 3% glycerol, 1 mM DTT, and 150 mM KCl). The detailed purification procedure can be found in SI. After affinity purification, the protein was further purified with gel filtration chromatography. The concentrated protein was loaded onto Superdex 200 column (GE Healthcare) coupled to an Akta FPLC purifier (Cytiva Life Science). The peak fractions were collected and analyzed with SDS-PAGE.

### Enzyme solution preparation

Prepare 5% (w/v) hemi-cellulase enzyme solution containing 0.2 M mannitol, 20 mM MES (pH 5.7) and 20 mM KCl. Incubate the solution in water bath at 55°C for 10 min to inactivate proteases and enhance enzyme solubility. Cool to room temperature (25°C) and add 0.1% BSA and 10 mM CaCl_2_. The enzyme solution should be prepared fresh.

### Plant seedling preparation

Sterilize the Arabidopsis seeds with 70% ethanol for 15 min and then 100% ethanol for 5 min. Air-dry the seeds in a laminar flow hood for 30 min and sow the seeds in a sterile plate containing autoclaved ¼ MS liquid medium supplemented with no sugar or DI water. Expose the seeds to 48 h of daylight or far-red light after overnight daylight exposure (12-15 h) at 25°C.

### Partial enzymatic digestion of cell wall and peptide treatment

Incubate the seedlings in enzyme solution for 4 h in dark and wash three times with rinsing buffer containing 0.2 M Mannitol, 4 mM MES (pH 5.7) and 15 mM MgCl_2_. Dry the seedlings on filter paper and submerge in protein solution (1 mg/mL in HEPES buffer). Incubate the seedlings for 12 h in dark at 25°C. The seedlings were rinsed with distilled water and then viewed using a Leica Confocal SP8-SMD microscope with a laser excitation wavelength of 488 nm.

## RESULTS AND DISCUSSION

As a proof of concept, we used GFP flanked by two NLS motifs (NLS-GFP-NLS-His) to assess the functionality of PECWD to detect successful nuclear GFP delivery. We utilized 3-day-old wildtype Arabidopsis seedlings due to their developing immature cell walls when compared to the more rigid structure of the mature cell walls of older seedlings or adult plants. Moreover, the miniature dimensions can accommodate confocal imaging and address the need for scaling up. The absence of sugar in the growth medium was useful not only for inhibiting the growth of fungus, but also for minimizing the carbon source that is required for biosynthesis of cell wall polysaccharides. Seedlings pretreated with 5% hemi-cellulase enzyme for 4 h and then incubated with 1 mg/mL NLS-GFP-NLS-His solution for 12 h were imaged using a confocal laser microscope (Leica SP8-SMD) and intense nuclear GFP fluorescence signals (round green nucleus) can be detected from the epidermal cell layers of cone-shaped root cap (Fig 2a). Without enzyme pre-treatment (Fig 2b) or protein incubation (Fig 2c), no nuclear GFP fluorescence signal was observed. The timing of 4 h enzyme pre-treatment was determined based on standard protocol for preparing Arabidopsis mesophyll protoplasts. As another piece of supporting evidence for NLS activity: while some cytosolic GFP localizations were observed, no GFP signals were found in the nucleus (Fig S1). It is worth noting that no positive nuclear GFP localization was detected in other tissues including root elongation zone, hypocotyl and cotyledon (Fig S2), presumably due to defective cell wall biosynthesis on fast-dividing root cap cells.^14^ In addition to hemi-cellulase, we also explored the potential of other cell wall-degrading enzymes that can serve as candidates for PECWD. While the use of Pectinase and Macerozyme results in the removal of root cap (Fig S3), no nuclear GFP fluorescence was detected on seedlings pretreated with cutinase (Fig S4). Together these results support our hypothesis that partially digested crude cell walls have enhanced permeability to cargo proteins.

**Figure 2.**
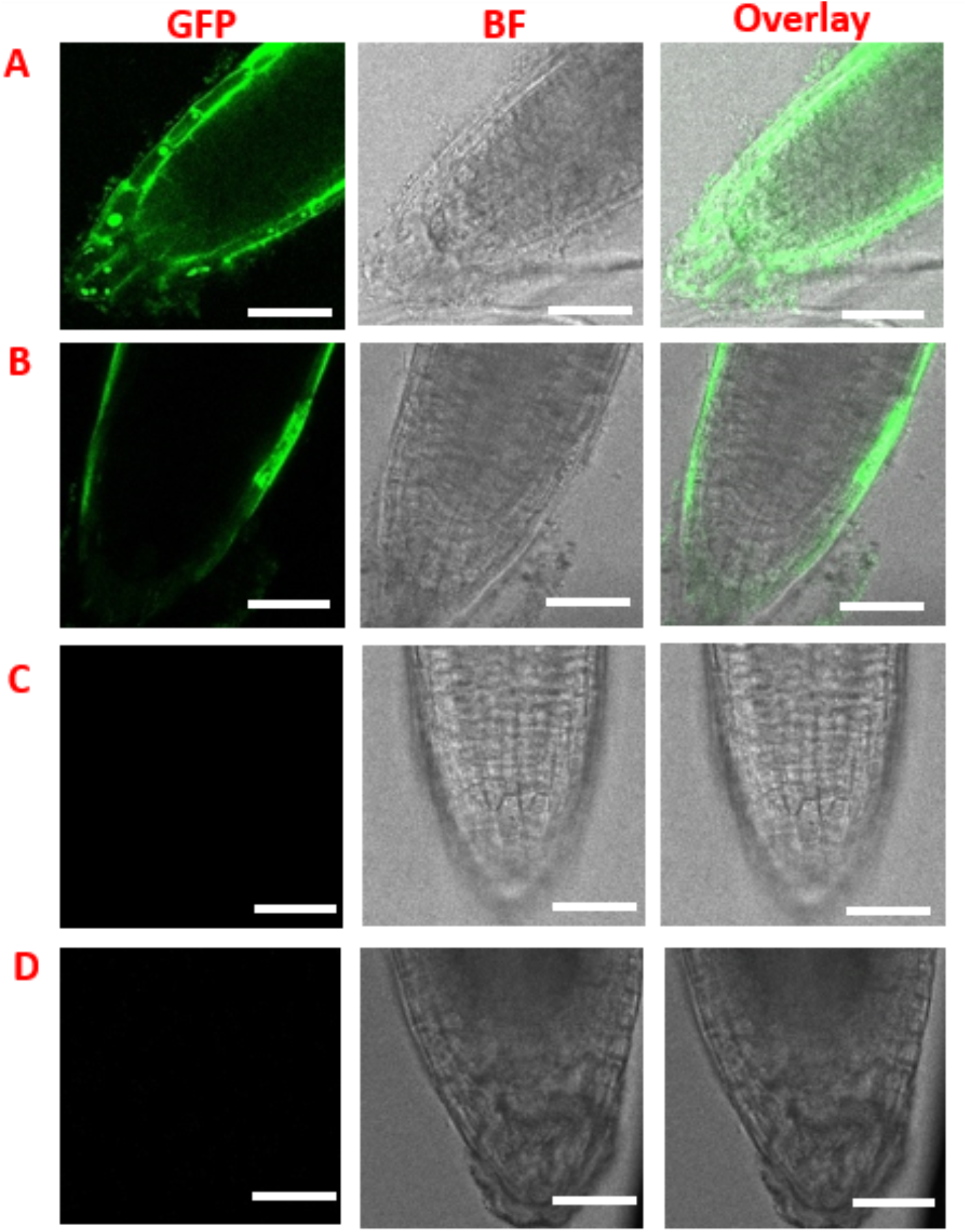
Nuclear internalization of GFP peptide in light-grown Arabidopsis seedlings. (A) Hemi-cellulase digested seedlings incubated with 1 mg/mL NLS-GFP-NLS-His for 12 h. (B) Intact seedlings incubated with 1 mg/mL NLS-GFP-NLS-His. (C) Hemi-cellulase digested seedlings without peptide incubation. (D) Intact seedlings without enzyme digestion and peptide incubation. The scale bars are 40 µm.

After validating PECWD for quantitative protein delivery on epidermal cell layers, we next assessed whether our approach could be further exploited to deliver the protein into deeper cell layers where meristematic cells are embedded. The biosynthesis of cell wall polysaccharides is fueled by carbon fixed by solar energy during photosynthesis. We hypothesize that photosynthesis can be inhibited upon switching from daylight to far-red light,^15^ resulting in a more defective cell wall structure that allows proteins to diffuse into deeper cell layers. After incubating hemi-cellulase digested far-red grown Arabidopsis seedlings with 1 mg/mL NLS-GFP-NLS-His solution for 12 h, both confocal (Fig 3a) and 3D images reconstructed by the Z-stack images (Fig S5) revealed that nuclear GFP fluorescence signals were detected throughout all cell layers of cone-shaped root cap. Similar to the light-grown Arabidopsis seedling, far-red grown seedlings, when not pretreated with hemi-cellulase or incubated with protein solution showed no nuclear GFP signals (Fig 3b and 3c). These results align with previous findings that far-red light exposure can alter morphological and physiological properties of Arabidopsis root.^15-16^ PECWD, when coupled with biological modifications of plant tissues, may offer a new approach to engineer meristematic cells and generate stably transformed plants in a culture-free manner.^2^

**Figure 3.**
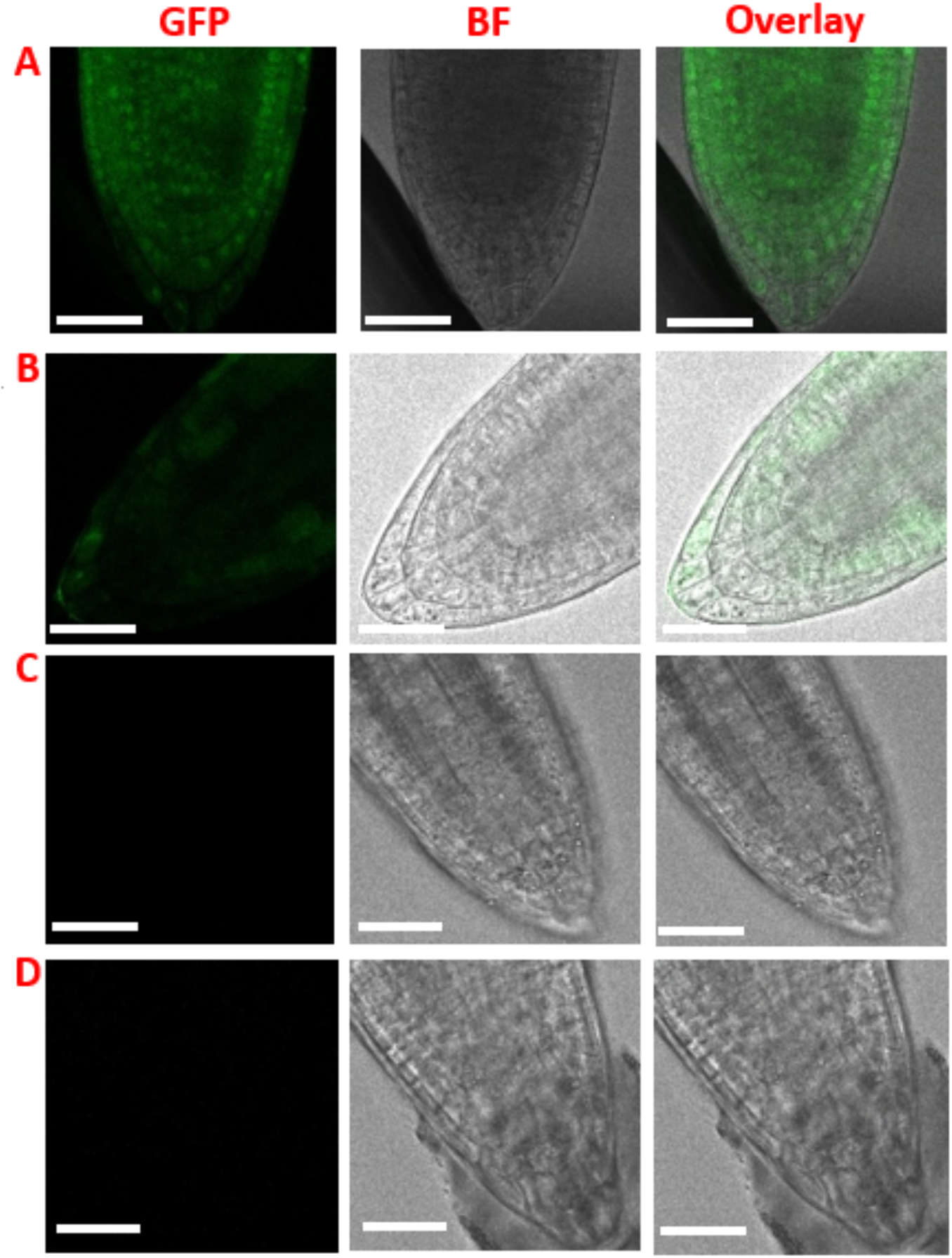
Nuclear internalization of GFP peptide in far-red grown Arabidopsis seedlings. (A) Hemi-cellulase digested seedlings incubated with 1 mg/mL NLS-GFP-NLS-His for 12 h. (B) Intact seedlings incubated with 1 mg/mL NLS-GFP-NLS-His. (C) Hemi-cellulase digested seedlings without peptide incubation. (D) Intact seedlings without enzyme digestion and peptide incubation. The scale bars are 100 µm.

While the above strategy is useful, the delivery of larger proteins is required to facilitate the major goals of plant delivery such as gene editing. This motivated our design of a Cas9 version of recombinant protein containing a GFP-tagged Cas9 flanked by two NLS, with a molecular weight of 188 kDa almost seven times as large as GFP. As indicated in Fig 4a, incubation with GFP-tagged Cas9 solution results in no GFP localization, whereas intensive GFP fluorescence was detected in Arabidopsis protoplasts with fully digested cell wall prepared by a standard procedure (Fig 4b).^2, 12^ It should be noted that longer digestion times and higher enzyme concentrations did not result in the delivery of Cas9 protein into the plant cell (Fig S6).

**Figure 4.**
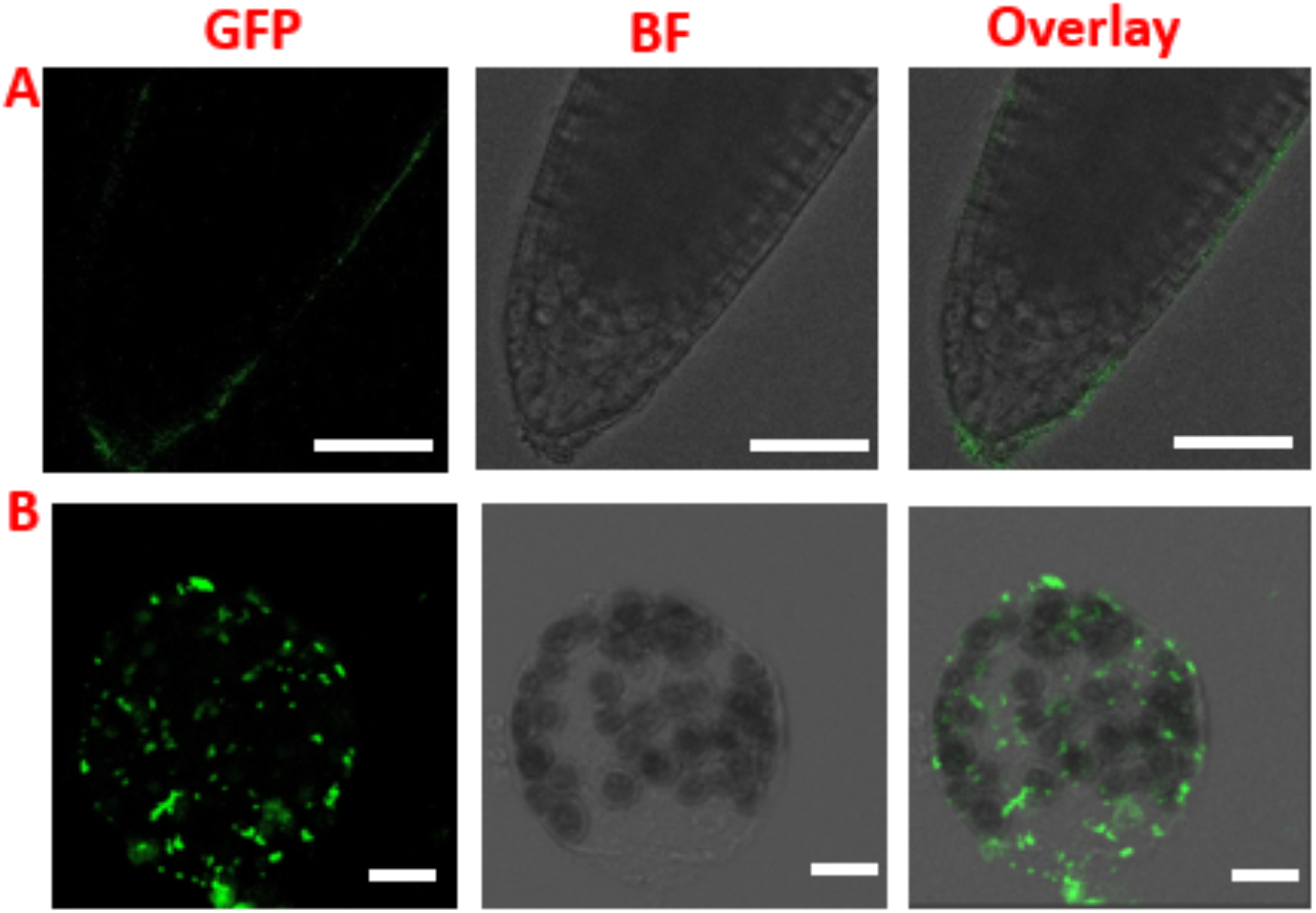
Nuclear internalization of GFP-tagged Cas9 peptide in far-red grown Arabidopsis seedlings. (A) Hemi-cellulase digested seedlings incubated with 1 mg/mL GFP-tagged Cas9 solution. The scale bars are 40 µm. (B) Arabidopsis protoplast incubated with 1 mg/mL GFP-tagged Cas9 solution. The scale bars are 10 µm.

Our study provides the first piece of evidence that partial digestion of the plant cell wall enables direct protein delivery into plant cells. However, our findings are a departure from the prevailing assumption that efficient delivery of proteins only occurs when the cell wall is completely removed. Two factors could be responsible for the discrepancy. As existing plant transformation methods mostly rely on protoplast culture (PEG, liposome, and electroporation) or physical disruption of the cell wall (particle bombardment, microinjection, and needle puncture), methods involving an “intermediate state” at which the cell wall is partially degraded have yet to be explored. Our PECWD represents an ideal way to explore the degree to which this “intermediate state” can be tuned to improve protein delivery efficiency. In addition, the majority of emerging cell penetrating peptide (CPP)^9^ and nanoparticles (NP)^11^-based techniques often suffer from low reproducibility. While the exact origins of these variabilities remain unclear, they are fundamentally connected to the structure of the plant cell wall and the resulting complex interactions that influence intercellular and intracellular protein translocation. The use of enzymatic pretreatment may help to overcome the obstacle of low efficiency and reproducibility associated with CPP or NP-mediated uptake of proteins.

The concept of utilizing a partial enzymatic cell wall digestion (PECWD) may be combined with various type of nanomaterials,^7,17^ expanding the utility of nanotechnology beyond the plant cell wall barrier and the current stage of proof-of-concept.^11, 18^ In addition to a model dicot plant Arabidopsis thaliana, our approach of partially disrupting cell wall polysaccharide may be applied to other plant species. The composition and structure of plant cell walls can differ among cell types and development stages, necessitating optimizations of enzyme type, enzyme concentration, and digestion time on a case-by-case manner. Ultimately, a molecular level understanding of the structure-function relationship between the cell wall and the permeability will help to unravel the origin of the significant variabilities observed in many biomolecule delivery methods, paving the way toward rational engineering of plants.

## ASSOCIATED CONTENT

### Supporting Information

**The Supporting Information is available free of charge at**

## AUTHOR INFORMATION

### Notes

The authors declare no competing financial interests.

## ACKNOWLEDGMENT

Q.G. and Z.W. acknowledge the support from Carnegie Science Venture Grant.

