## Supplemental Information for "Direct Nuclear Delivery of Proteins on Living Plant via Partial Enzymatic Cell Wall Digestion"

### Partial Enzymatic Relaxation of Plant Cell Wall Enables Nuclear Delivery of Protein into Deep Cell Layers

**This PDF file includes:**

Tables
Figures S1 to S8
SI References

***Recombinant protein purification.*** Transform the pET28a plasmid carrying coding sequence into BL21(DE3) *E. coli* strains for protein expression. Culture a single transformed clone in 10 mL LB medium overnight. Transfer the culture into 2L LB medium for large-scale induction. Grow *E. coli* to $OD_{600}$ = 0.8 at 37℃ with shaking and cool to 16℃ before induction. Incubate protein induced with 300 μM isopropyl-β-D-thiogalactoside (IPTG) overnight. Upon centrifugation, resuspend the bacterial pellet in HEPES buffer (20mM HEPES, 150 mM KCl, 3% glycerol, pH 7.5). Lyse the pellet with ultrasonication and purify the protein with $Ni^{2+}$ columns (Invitrogen, R901-01), remove non-specific proteins by washing the column with 20 mM imidazole in HEPES buffer. Elute protein from the column with 300 mM imidazole in HEPES buffer. After affinity purification, purify the protein with gel filtration chromatography. Load 500 μL concentrated protein onto Superdex 200 column (GE Healthcare) coupled to an Akta FPLC purifier (Cytiva Life Science). Collect the peak fractions and analyze with SDS-PAGE. Aliquot the fraction and flash-frozen in liquid nitrogen and store in a cryo-freezer (-80℃).[1]

***Confocal imaging.*** Confocal imaging was performed with a Leica TSC SP8 point scanning confocal microscope equipped with a white light laser (WLL) exciting at 488 nm. The emission for every pixel was detected with a single-channel PMT detector. The range was set between 498 nm to 530 nm using an acousto-optical tunable filter (AOTF). The image resolution was 0.24 um in X and Y and 1 um in Z. The scanning speed was set to 400 lines per second. Brightfield images were acquired by a different PMT detector measuring 488 nm transmitted light.

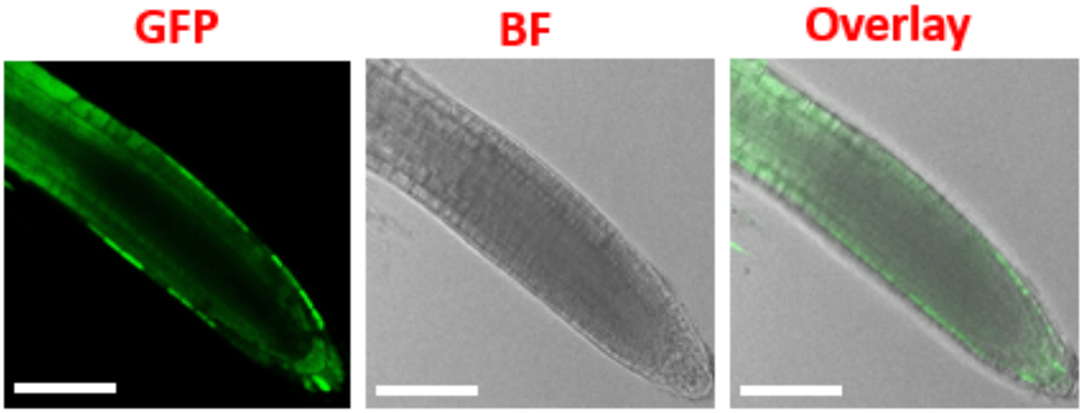


**Figure S1**. Hemi-cellulase digested daylight grown Arabidopsis seedlings incubated with 1 mg/mL GFP-His solution. The scale bars in insets are 50 μm.

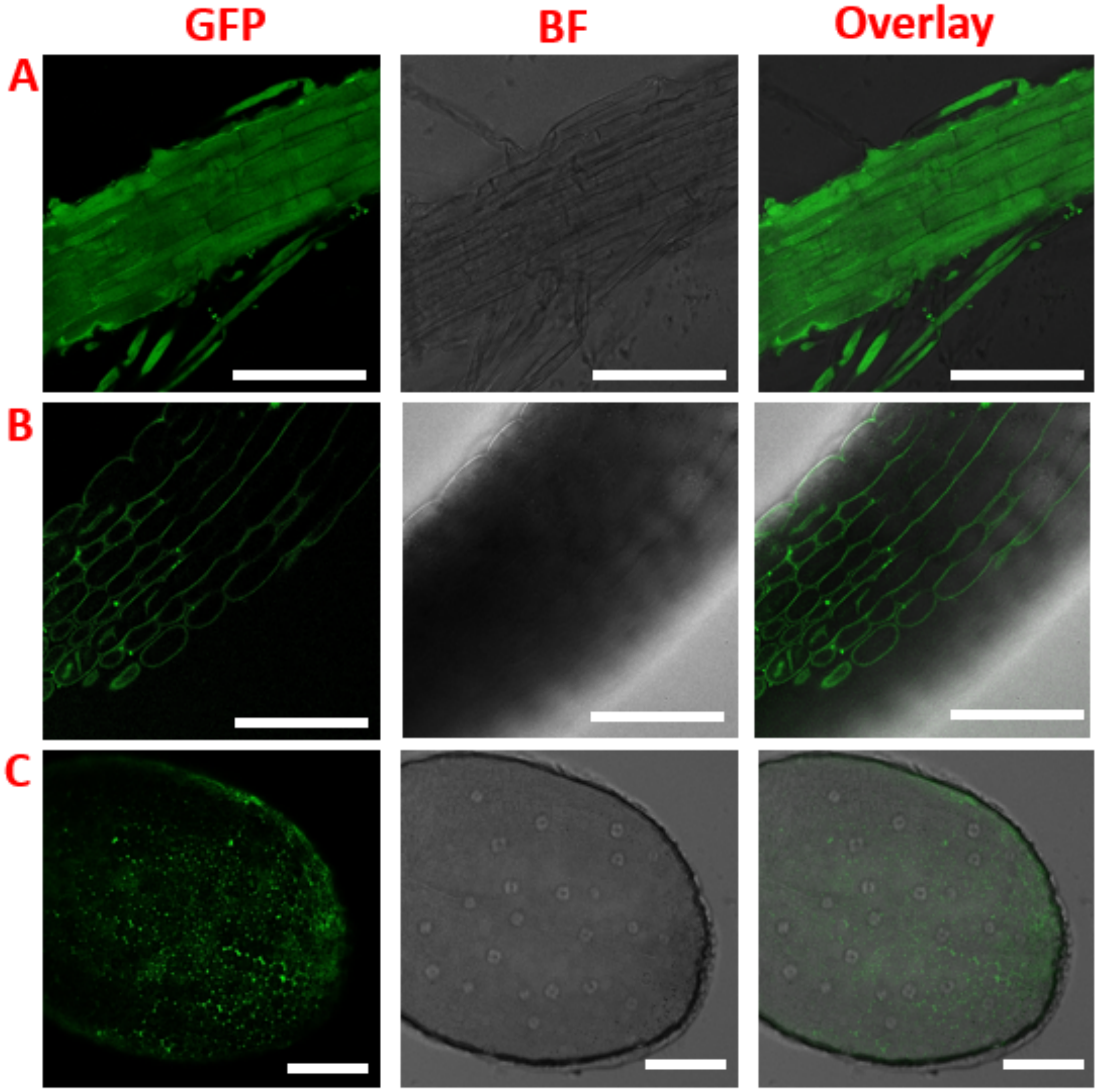


**Figure S2.** Hemi-cellulase digested Arabidopsis seedlings incubated with 1 mg/mL NLS-GFP-NLS-His solution. Representative confocal images of (A) root elongation zone, (B) hypocotyl and (C) cotyledon. The scale bars are 100 μm.

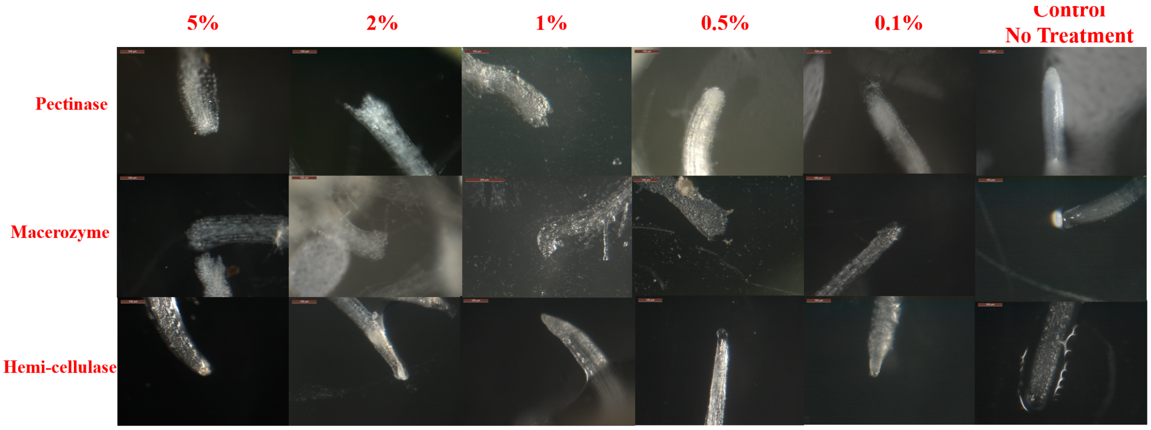


**Figure S3.** Far-red grown Arabidopsis seedlings incubated with (A) Pectinase, (B) Macerozyme and (C) Hemi-cellulase at different enzyme concentrations.

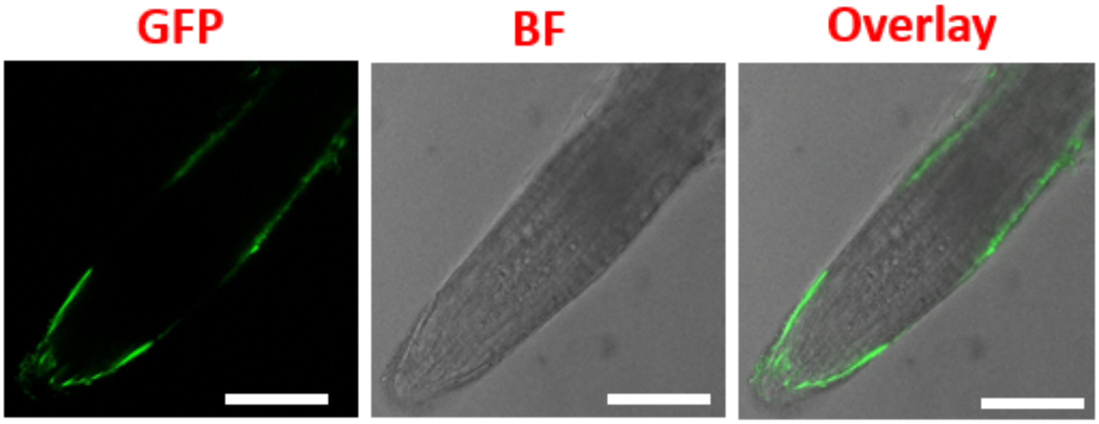


**Figure S4.** Cutinase digested Arabidopsis seedlings incubated with 1 mg/mL NLS-GFP-NLS-His solution**.** The scale bars are 50 µm.

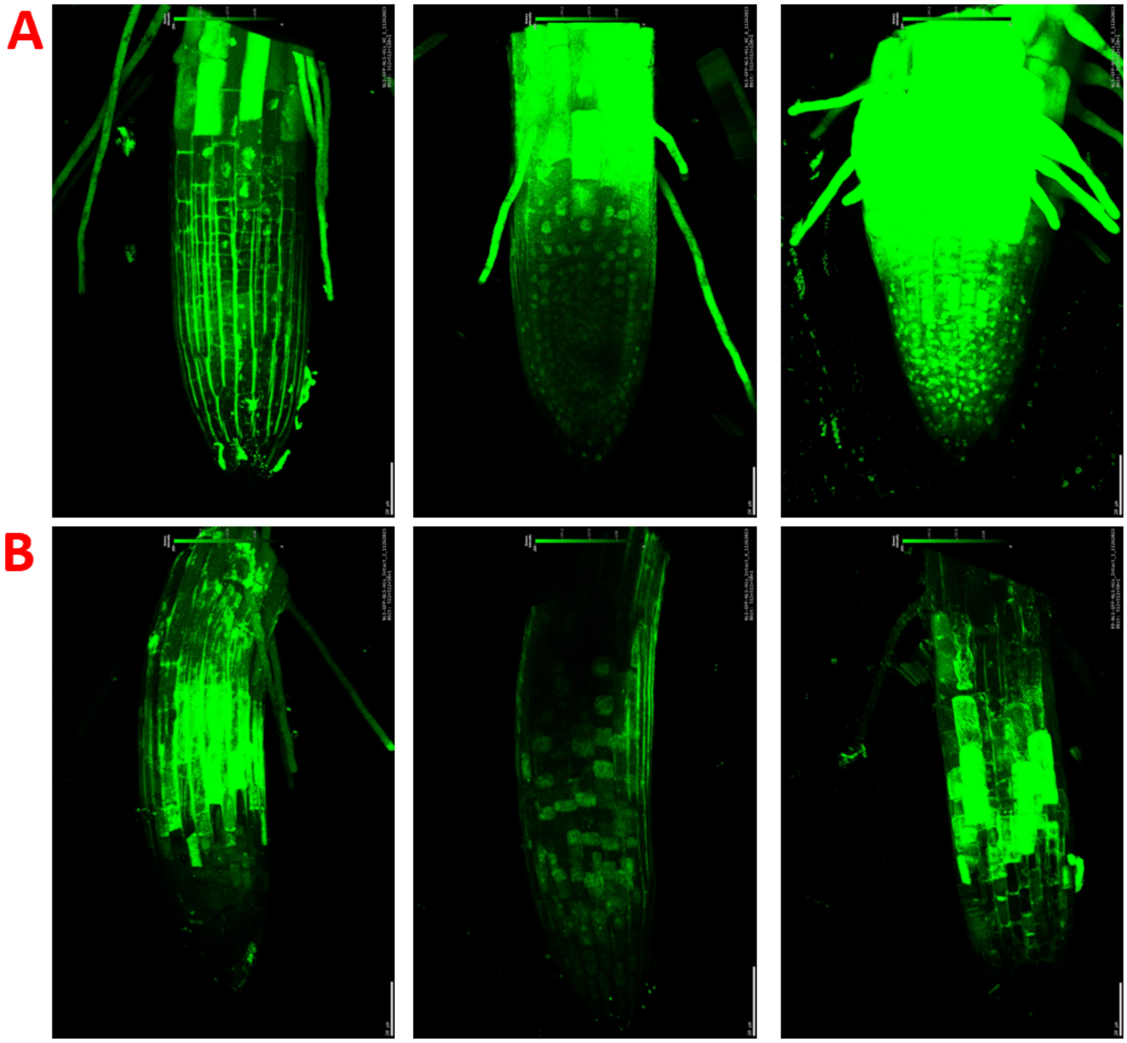


**Figure S5.** Nuclear internalization of GFP peptide in far-red grown Arabidopsis seedlings. 3D images reconstructed by the Z-stack images (A) Hemi-cellulase digested seedlings incubated with 1 mg/mL NLS-GFP-NLS-His for 12 h**.** (B) Intact seedlings incubated with 1 mg/mL NLS-GFP-NLS-His for 12 h.

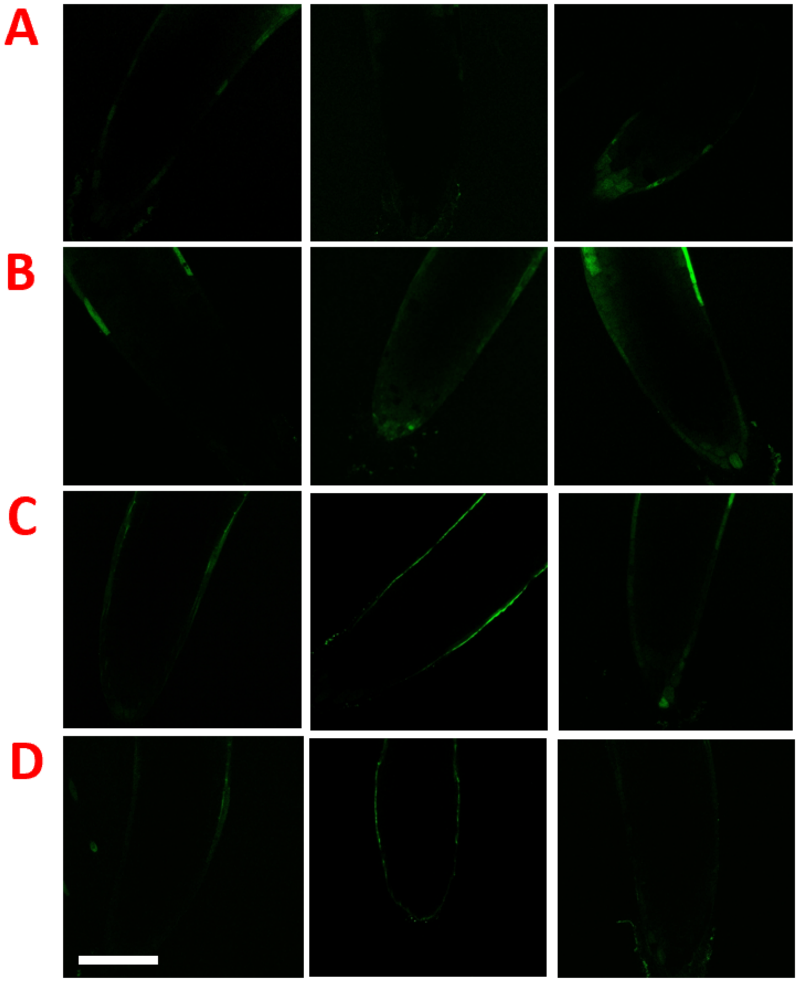


**Figure S6.** Hemi-cellulase digested Arabidopsis seedlings incubated with 1 mg/mL NLS-Cas9-NLS-GFP-Motif-His solution. Representative confocal images of far-red grown Arabidopsis seedlings digested by (A) 0%, (B) 5%, (C) 10%, (D) 20% Hemi-cellulase for 6 h (left panel), 12 h (middle panel) and 24 h (right panel).

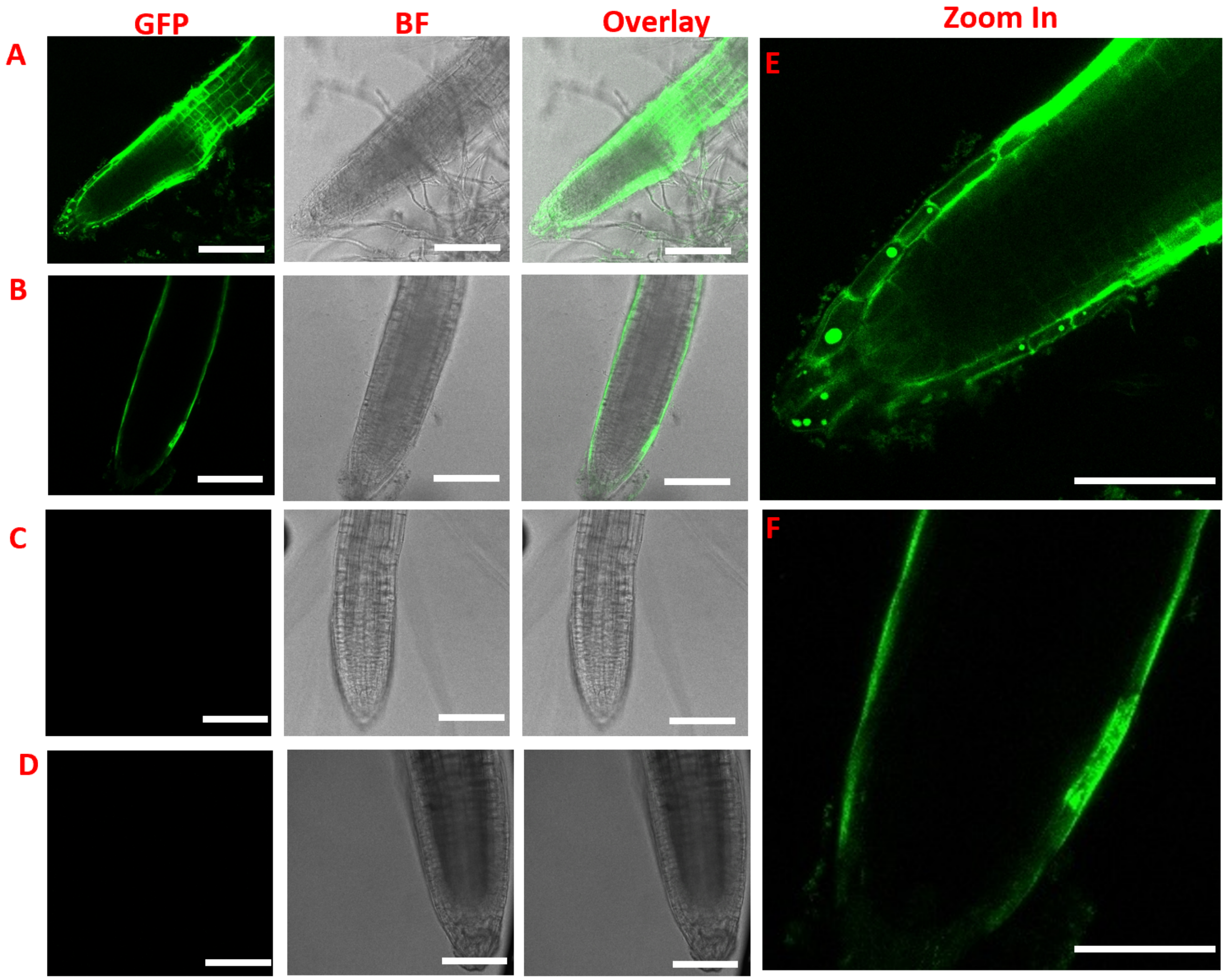


**Figure S7. Nuclear internalization of GFP peptide in light-grown Arabidopsis seedlings.** (A) Hemi-cellulase digested seedlings incubated with 1 mg/mL NLS-GFP-NLS-His for 12 h. (B) Intact seedlings incubated with 1 mg/mL NLS-GFP-NLS-His. (C) Hemi-cellulase digested seedlings without peptide incubation. The scale bars are 100 µm. (D) Intact seedlings without enzyme digestion and peptide incubation. (E) Zoom-in from panel A showing nuclear GFP signals in root cap as the result of successful delivery. (F) Zoom-in from panel B showing cytosolic GFP signals only. The scale bars are 50 µm.

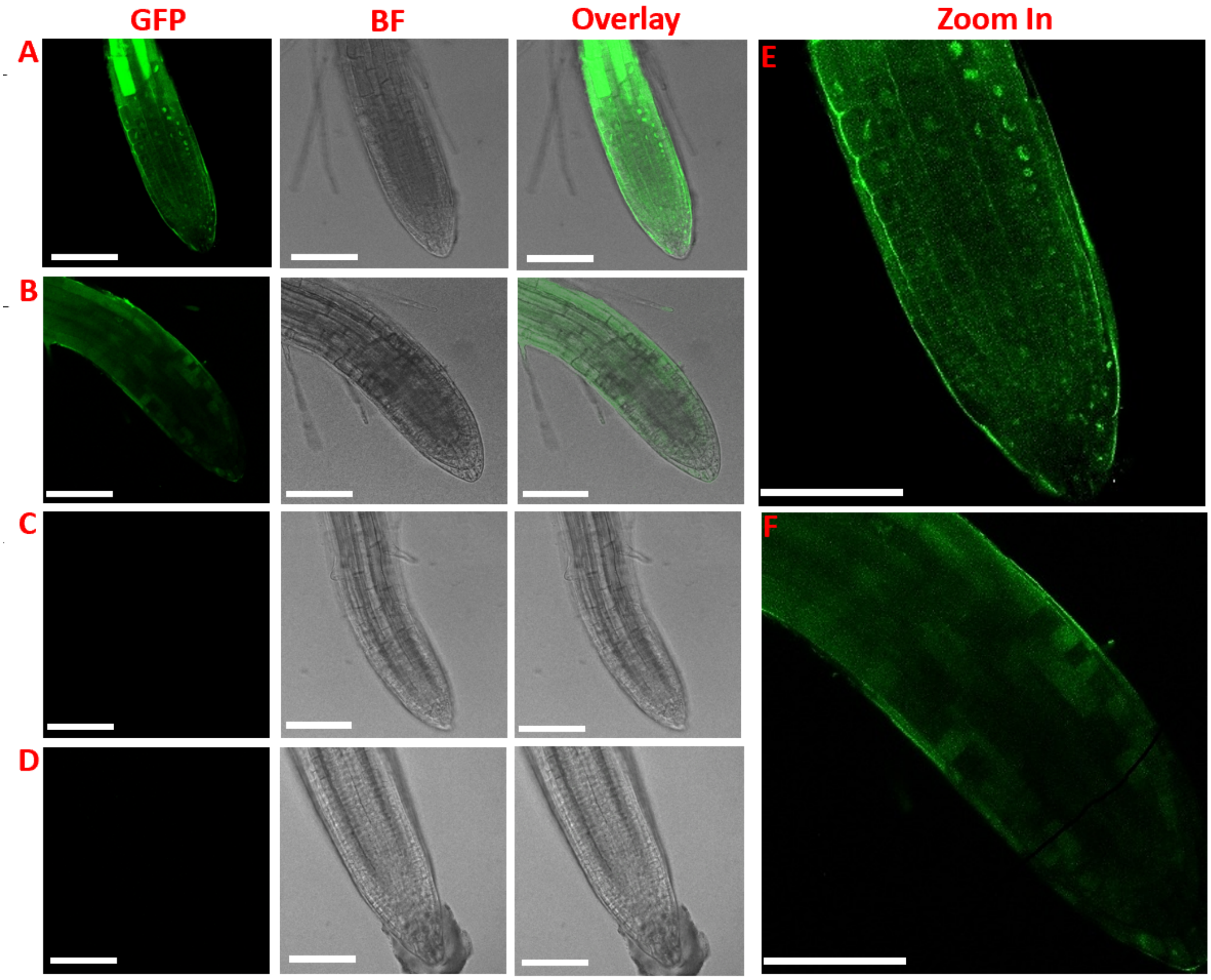


**Figure S8. Nuclear internalization of GFP peptide in far-red grown Arabidopsis seedlings.** (A) Hemi-cellulase digested seedlings incubated with 1 mg/mL NLS-GFP-NLS-His for 12 h. (B) Intact seedlings incubated with 1 mg/mL NLS-GFP-NLS-His. (C) Hemi-cellulase digested seedlings without peptide incubation. The scale bars are 100 µm. (D) Intact seedlings without enzyme digestion and peptide incubation. (E) Zoom-in from panel A showing nuclear GFP signals in root cap as the result of successful delivery. (F) Zoom-in from panel B showing cytosolic GFP signals only. The scale bars are 50 µm.
